# Inclusive medical rehabilitation for persons with disability due to leprosy, lymphatic filariasis, and diabetes mellitus: Mapping the gap in three leprosy endemic districts in Indonesia

**DOI:** 10.1101/511097

**Authors:** Hanifa Maher Denny, Yudhy Dharmawan, Praba Ginandjar, Bagoes Widjanarko, Ayun Sriatmi, Aizzatul Umamah, Mateus Sakundarno, Widya Prasetyanti

## Abstract

Medical rehabilitation for person with disability in Indonesia is still an issue. This research aimed to explore Inclusive medical rehabilitation for persons with disability due to leprosy, lymphatic filariasis, and diabetes mellitus in three regions in Indonesia. The qualitative study was employed to gather data from disability patients, health workers in PHCs, medical rehabilitation services for leprosy, DM, and LF in hospitals.

The results indicated that the gap on medical rehabilitation for person with disability due to leprosy, lymphatic filariasis and diabetes mellitus in three regions were due to some differences in their geographical aspects, availability of referral hospital for treating leprosy and filariasis, supervision, human resource competencies.

## Introduction

Medical rehabilitation in Indonesia remains limited in terms of availability and accessibility to the majority of people needing it [1,2]. There are medical rehabilitation centres in all major cities, and even district hospitals must offer basic rehabilitation services. This however does not mean that all the relevant services are obtainable nationwide, that the professional staff is available, or that patients or persons with disabilities are adequately informed about their options. For persons with disability living in poverty, lack of funds or information about their rights may cause them to go without any rehabilitative interventions their whole life [3].

The right to receive health services contains the elements of availability, accessibility, acceptability, and appropriateness (or quality) [2]. In Indonesia, people with leprosy often experience barriers in obtaining rehabilitation services [4]. For example, a previous study described barriers to medical rehabilitation services for persons with disability due to leprosy [5]. There are seven leprosy hospitals available throughout Indonesia, but only one offers the most relevant medical rehabilitation (e.g. surgery, prosthetics & orthotics, physiotherapy, occupational therapy, including here also guidance to home-based self-care) [3]. Long distance to reach the nearest leprosy hospital limits the access for most leprosy-affected persons. Furthermore, access to general hospitals is often restricted by stigmatising and discriminating attitudes of hospital staff [4]. Furthermore, information accessibility is constrained for many people in remoter areas and with low level of education [6].

Many persons affected by leprosy are not members of any insurance scheme as they do not have valid identity cards [7]. Additionally, economical access or affordability is often not ensured when people have to stay for prolonged times in leprosy hospitals waiting for services, or have to bring family members to care for them, both reducing family income [8]. Acceptability of medical services is low in some leprosy hospitals where facilities and services hardly respect human dignity, as clearly evident when visiting these hospitals. Appropriateness or quality for services differs and may only benefit those who have the resources.

Other diseases such as lymphatic filariasis (LF) and diabetes mellitus (DM) can cause disabilities similar to leprosy in several aspects [9–11]. All require care and self-care to deal with chronic impairments especially to lower limbs [12–14]. Both in leprosy and DM, peripheral neuropathy is common, which may lead to wounds when pain is not felt, and ultimately to amputations [9,11]. Simple interventions such as wearing protective footwear may prevent worsening of impairments for people with insensitive feet caused by neuropathy, while tendon transfers in leprosy restore function and mobility [12]. On the other hand, people with LF develop lymphedema on their limbs, which could be reduced and kept from worsening through home-based lymphedema management [13]. In later stages surgery might be beneficial [15].

Medical rehabilitation plays an important role in all three diseases [16,17]. For the diseases, self-care is essential, which ideally should be taught by hospitals and primary health cares (PHCs) [12–14]. Thus, it is important to develop models used for advocacy and implementation of inclusive medical rehabilitation for persons affected by leprosy, LF and DM. The aim of this study is to generate knowledge on the gap of medical rehabilitation services in three leprosy endemic areas, in order to improve comprehensive rehabilitation services.

## Materials and Methods

### Study design

This was a qualitative study designed as baseline data to map the gap of inclusive medical rehabilitation for leprosy, LF, and DM. The aim of this study is to systematically and comprehensively collect information regarding the enabling and disabling factors influencing the successful provision of inclusive medical rehabilitation services for persons with disability due to leprosy, LF and DM.

### Study site

Selection of the study site was based on endemicity of leprosy according to data from Ministry of Health. The study was carried out in three leprosy districts, namely: Pekalongan District, PALI District, Bima District (leprosy prevalence was 1.33 %, 1.73%, and 2.2 % respectively).

### Informant

The informants of this study were disability patients, health workers in PHCs, medical rehabilitation services for leprosy, DM, and LF in hospitals. The informants were selected purposively in Penukal Abab Lematang Ilir (PALI) District (Sumatera Selatan Province), Pekalongan District (Jawa Tengah Province), and Bima District (Nusa Tenggara Barat Province). The informant of this study consisted of three groups: 29 patients, 41 health workers, and 17 experts.

### Conceptual Research Framework and Variables

The conceptual framework implemented the Innovative Care for Chronic Condition (ICCC) approach. The framework was initialy published by World Health Organization (WHO) as an innovative method to manage current care and prevent future chronic condition [18]. The approach consists of 3 dimensions: from the point of view of health services, community, and patients and their families. All the three dimensions were obtained from DM, leprosy, and LF patients.

### Data Collecting Technique

Data was collected by indepth interview. Instruments for data collection consisted of questionnaire and observation list. The instruments were tested in a district that has similar characteristics with the study locations.

### Data Analysis

Data was analysed by content analysis according to the response from the informants, which then was attributed to the context of variables. The data was analyzed to describe the gap between each variable of the three programs (LF, DM, and Leprosy) and also the three locations of study.

### Ethics Statement

Ethical approval was issued by the Committee of Public Health Research Ethics, Diponegoro University (233/EC/FKM/2016). Writtien informed consent was acquired from all subjects. For subject under 18 years old, informed consent was provided by a parent on behalf of the subject.

## Result

### Characteristics of informants (patients, health workers, experts)

The number of patients in Pekalongan, PALI, and Bima was 29 respondents (10, 11, and 8 respectively), consisted of 16 females and 13 males. The average age was 51 years old (ranged 14-72 years old). The patients had several types of occupation but mostly were housewives (27.6%), followed by farmer (13.8%). Sixty seven point nine percent of patients were married while 37.0% did not graduate from elementary school. The duration of illness was 2-15 years for Leprosy, 5 years for DM, and 32 years for LF.

The mean age of health workers was 39 years old, and the majority of them were female. The majority of health workers were graduated from Diploma of Nursing (44.7%), married (92.1%), and were the native of the district. The average distance from the residence of health worker respondents to their workplace is 4.6 km with the closest distance 15 meters and the longest distance 25 km. 88.6% of respondents ride motorcycle as transportation to the workplace. Respondents have worked in their respective workplace on average of 12 years.

In this study, experts consisted of physiotherapist (22.2%), internist (22.2%), dermatologist (11.1%), surgeon (22.2%), general practitioner (11.1%), and professional nurses (11.1%). The majority of experts were male (55.6%). The average age of respondents is 47 years old with the youngest is 45 years old and the oldest is 51 years old. 4 of 9 respondents are known to have studied the disease for an average of 15 years, with the shortest time to explore the field of disease is 2 years and the longest is 24 years.

### The Gap of Health Services for Leprosy, Diabetes Mellitus, and Lymphatic Filariasis

#### Infrastructures

Leprosy, DM, and LF patients reported similar statement about the PHC infrastructure for which the patients are sufficiently satisfied with the building condition and the facilities of health services. The positive response was reported by patients in Pekalongan and Bima Districts that the facilities are sufficient and helpful. In a similar comment, a patient from outside Java stated the facility in health service was convenient. Other patients felt grateful for the services.

Health workers reported that the building condition had been a major concern for the health management, which was addressed by continuous effort to improve the condition. However, there were occassions when examinations were conducted where the health workers were on duty, for example, if the health worker is responsible for treating the leprosy and also TB, then the examination for both diseases will be conducted in the same room. It was also known that the total building area in the PHCs affected the availability of examination rooms. With regards to the temporary room for DM patients, one health worker addressed the importance of renovating the health facility. A health worker informed that while the medical rehabilitation for LF in Pekalongan District was not yet available, the mass drug administration had been launched. The annual launching is scheduled in October. She also added that the ideal facility for combining the treatment of Leprosy, LF and DM should be in the form of a rehabilitation room. Rehabilitaion for TB and Leprosy patients should be isolated in a special room and the care givers assigned should also be separated for each disease.

#### Inputs and Equipment

Health workers stated that connection to the sewage in the PHCs is available along with the water system. Data regarding the availability of basic equipment for the diagnosis and therapeutic in the PHC facilities was obtained from a checklist filled out by the health workers. The results can be seen in the following table.

The availability of basic equipment for the diagnosis and therapeutic is presented in Table 1. It can be seen that the availability of basic equipment for the LF diagnosis and therapeutic is the most incomplete when compared to others. In detail, the average of the availability of basic equipment for LF diagnosis and therapeutic is only 38.5%, creating a 21.5% gap with Leprosy and 38.8% with DM. Based on district, it can be seen that Pekalongan has more equipment, although only a slight gap was found between three district. In detail, the average of the equipment availability in Pekalongan is 68.7%, creating a small gap of 7.7% with Bima and 15% with PALI. In contrast to the health worker, patients described how the equipment always available and there were no signs of deficiency. In contrast to the data given by the health workers, patients described how the equipment was always available and that there were no perceived signs of a deficiency. Other patient also informed that the availability of the service material was complete.

**Table 1.** Percentage of availability of the basic equipment for the diagnosis and therapeutic

| District | Leprosy | DM | LF | Average |
| --- | --- | --- | --- | --- |
| Pekalongan | 76 | 95 | 35 | 68.7 |
| PALI | 52 | 67 | 42 | 53.7 |
| Bima | 52 | 70 | n/a | 61.0 |
| Average | 60 | 77.3 | 38.5 |  |

In terms of laboratory availability for the clinical checkup, all PHCs observed in Pekalongan have a laboratory despite its lack of equipment. Meanwhile, both in PALI dan Bima Districts, patients acknowledged that the laboratory function in the PHCs was not optimal. Leprosy patients in Bima District, are therefore required to use the service of private laboratory which sometimes involves considerable amount of money. In addition, in all the three study sites, each healt centre has at least one ambulance to help services. Most patients were also aware that there was ambulance available in each PHC.

#### Pharmaceutical

Data regarding the availability of generic and supporting drugs in the PHC was obtained from a checklist filled out by the health worker. Results can be seen in the following table.

**Table 2.** The percentage of availability of the generic and supporting drugs

| Districts | Leprosy | DM | LF | Average |
| --- | --- | --- | --- | --- |
| Pekalongan | 68 | 16.7 | 70 | 51.57 |
| PALI | 100 | 41.9 | 50 | 63.97 |
| Bima | 100 | 16.7 | n/a | 58.35 |
| Average | 89.3 | 25.1 | 60 |  |

The result indicated the availability of generic and supporting drugs for leprosy is more complete than DM and LF. However, it should be taken into account that some types of drugs were not available at PHC. There were significant differences between the percentages gap with LF (29.3%) and with DM (64.2%). It can also be seen that the drugs availability in PALI District were more complete than in Pekalongan and Bima Districts. However, there was only a small gap found between the three districts.

The Government of Indonesia is organizing programs to provide medicines for leprosy, DM, and LF covered by the BPJS Health program, in the form of health insurance. DM drugs are included in the *Prolanis Program*, associated with other non-communicable diseases. While the drug for LF patients only exist in PHC, therefore the hospitals often repatriate LF patients. This was the reason hospitals were not as knowledgeable in treating LF patients.

#### Human Resources

According to the patients, the presence of health workers serving the PHC patients is sufficient. The human resources in PHC and in the hospital are very different. In PHC, health workers are usually given the task of holding one or more programs in addition to their main program. For example in DM, the program is incorporated in the other non-infectious disease program. There are leprosy officers who are also responsible for tuberculosis programs, some are assigned concurrently with LF. This is due to the limited number of qualified health workers that can be given the responsibility. As a result, the program officers were not able to share the attention of both programs fairly and harmoniously.

On the other hand, health workers at referral hospitals such as Dr. Rehatta Hospital in Jepara-Central Java and Dr. Rivai Abdullah Hospital in Palembang-South Sumatra at least consist of a general practitioner, a dermatology specialist (for leprosy), an internist specialist (for DM), nurses, a physiotherapist, officers in orthotic prosthetic, and medical rehabilitation doctors. Unfortunately LF expert are not found in these two hospitals. However, an expert from Dr. Rivai Abdullah Hospital, who is a surgeon, stated that he was able to perform surgery for LF patients who already suffered swollen lymph nodes.

#### Supervision

Supervision for the implementation of the program of DM, leprosy, and LF in PHC is provided by the District Health Office (DHO). PHCs provide oral and written reports every three months to DHO. Unfortunately, not all officers are able to make quick and complete reports. This was stated by a health worker in Sape PHC (Bima District). In addition to reporting every three months, the PHC also holds internal monthly workshops for staffs aimed to increase the vigilance of the stakeholders of the other programs as well. It also reaffirms and indicates that, in fact, tracking and management of each disease is the responsibility of an individual officer.

#### Referral

Referral sites for leprosy rehabilitation exist only in Central Java and South Sumatra. Dr. Rehatta Hospital in Jepara, Central Java, became a reference only for Central Java Province, while Dr. Rivai Abdullah Hospital became a reference throughout Sumatra and West Kalimantan. Bima District, on the other hand, has no experience to refer leprosy patients to medical rehabilitation due to the long distance to the location of leprosy referral hospitals in Mataram or in Makassar.

Referrals for medical rehabilitation of DM in Central Java and South Sumatra may be targeted at local public hospitals in each district or in the same place as leprosy referral hospitals, since they receive services for medical rehabilitation with other causes such as stroke and traffic accidents. While in Bima District, there is no place for medical rehabilitation referral for DM as well as LF, and leprosy diseases. Referrals for medical rehabilitation due to LF have not been officially established in all three provinces, but an informant in South Sumatra stated that he may be able to handle the rehabilitation of LF patients.

## Discussion

The gaps can be seen in the three research areas (Pekalongan, Bima, and PALI Districts). The condition of health facilities and infrastructure in Central Java is relatively better than in South Sumatra and West Nusa Tenggara. Nevertheless, in areas directly adjacent to the Java Sea in Pekalongan District, many are experiencing floods [19], so the PHC is inundated by sea water forcing many health workers to wear boots. Therefore, several PHCs in Pekalongan District are rebuilt, having more than one floor. The condition of most PHCs in Bima District does not meet the standard/is not adequate, however there is one PHC in Sape with adequate building and facilities. On the other hand, seven PHCs in PALI District are acceptable.

According to the previous findings, it could be understood that the gaps on medical rehabilitation for leprosy, DM, and LF remain exist [19,20]. In this study, the gaps are mainly illustrated on self-care management and referral system. The LF and DM do not have self-care program, which is available for leprosy program. In LF and DM, the program only deal with passive finding and treatment only. Likewise, referral schemes for rehabilitation and advanced treatment are of no special nature as can be found in leprosy programs. Leprosy programs show special programs i.e. from active case finding such as contact surveys, RVS and school surveys.

Treatment of leprosy patients is also always accompanied by a self-care mechanism taught by the officer to the patient, in order to prevent the disease from worsening [21]. There is also a mechanism of rehabilitation through reconstruction, physiotherapy and the provision of aids (prosthesis) for people with disabilities level 1 or 2 [22]. DM program is managed in conjunction with other non-infectious diseases program, so there is no specificity of the DM program. If anyone is disabled by DM, there is no specific scheme for rehabilitation. This is also the case with LF, although it is a separate program in PHC the rehabilitation or self-care efforts do not become the focus in the treatment of patients. This self-care gap, as well as rehabilitation efforts, can actually be overcome by combining the care of disabled DM and LF patients into the treatment of leprosy patients which are currently underway [20]. In this case, medically self-care efforts and medical rehabilitation that have been applied to leprosy patients can also be applied to LF and DM patients.

This study indicated that in the region there are variations in leprosy referral system implementation inter-regional, especially from the aspect of availability of referral facilities and referral system. Refferal medical rehabilitation was only available in leprosy programs. The best referral system only exists in Java Island. In the research area in Central Java, they have referral hospital with a system to pick up patients who are ready to be rehabilitated by coordinating the Leprosy Supervisor in DHO and PHC. In South Sumatra, the leprosy referral hospital does not have a system such as in Central Java. RS is passively waiting for referrals from the region. In West Nusa Tenggara research area there is no clear referral mechanism nor any case of PHC referring leprosy patients to undergo rehabilition as there is no leprosy hospital in the area.

Regional policies greatly affect the implementation of medical rehabilitation services in the regions. National policy is required on the best referral mechanism for leprosy patients. Policy making also needs to be accompanied by the provision of facilities along with the division of working area, so that the reference facilities are easily accessible by the patient. It is also necessary to consider the infrastructures to the hospital, such as public transportation and good road conditions, for easy public access once the hospital is announced as able to treat disabled patients due to DM and LF. We suggest Leprosy Hospital may also use existing hospitals, with additional training for staffs. The hospital can also be equipped with therapists to assist the recovery of the patient. This policy should be executed considering the distribution of health services for rehabilitation should be implemented in all parts of Indonesia.

Limitation of this study such as information bias could be occurred in some translations of the specific indigenous dialect such as the name of the disease in the local.dialect. For example, "Ncola" in Bima refers to Leprosy. Therefore, the investigators asked the local health officer to make sure that the term of the disease in a local dialect was translated correctly.

## Conclusion

Gap of medical rehabilitation services remains exist between leprosy, diabetes, and filariasis. Medical rehabilitation program of leprosy is better than the other diseases. There was also a gap of medical rehabilitation services between Java and non-Java Islands.

1. Availability of facilities and referral system in Java Island was better. The worst services were found in east Indonesia (Nusa Tenggara Barat Province).
2. Availability of facilities, health officer conformity, and community acceptance are factors to minimalize stigma and are key factors of inclusive medical rehabilitation service.
3. Model of integrated medical rehabilitation between leprosy, filariasis, and diabetes may be developed by involving puskesmas as treatment and selfcare unit, and leprosy hospital as referal unit of reconstruction and rehabilitation.
4. Government should develop national policy and universal coverage for health insurance, which will be an operational reference in puskesmas and hospital level. Puskesmas, hospital and government should be oriented to community empowerment, with final goal to decrease leprosy stigma and increase community awareness on the importance of medical rehabilitation for leprosy, diabetes, and filariasis.

## Acknowledgments

The authors would like to thank all the participants of this study. The authors would also like to thank the corresponding Public Health Centers, Hospitals, and Distsric Health Offices.

## Supporting Information Legends

S1 Checklist: STROBE Checklist

